# Evaluating the solubility of compounds intended for intramammary infusion based upon tests conducted across a range of milk products using cephapirin sodium and benzathine as model drugs

**DOI:** 10.1101/2023.01.24.525429

**Authors:** Marilyn N. Martinez, Fang Zhao, David G. Longstaff, Justin J. Gabriel, Martin J. Coffey

**Affiliations:** US FDA Center for Veterinary Medicine, Office of New Animal Drug Evaluation, Rockville, MD; St. John Fisher University, Wegmans School of Pharmacy, Rochester, NY; Research and Development, Bausch & Lomb, Rochester, NY

## Abstract

The ability to evaluate drug solubility in milk and milk-related products has relevance both to human and veterinary medicine. Model compounds explored in a previous investigation focused on drug solubility assessments when delivered in milk-associated vehicles for administration to human patients. In the current investigation, we focus on the solubility of drugs intended for delivery via intramammary infusion to cattle. Because there are logistic challenges typically associated with obtaining raw milk samples for these tests, there is a need to determine potential alternative media as a substitute for raw bovine milk. Given the complexity of the milk matrix, aqueous media do not reflect the range of factors that could impact these solubility assessments. This led to the current effort to explore the magnitude of differences that might occur when substituting raw bovine milk with off-the-shelf milk products such as whole milk, skim milk, or reconstituted whole milk powder. We considered conclusions based upon the solubility assessments derived from the use of the model compounds studied in our previous report and compared them to conclusions obtained when testing two drugs with differing physicochemical characteristics that are approved for administration via bovine intramammary infusion: cephapirin benzathine and cephapirin sodium. Based upon these results, we recommend that whole milk or reconstituted whole milk can substitute for bovine raw milk for the solubility assessment of compounds intended for administration via intramammary infusion. However, unlike the human drug situation, these tests should be conducted at 38°C.

## Introduction

Bovine mastitis is a major health problem encountered both within the US and around the world [1]. Its importance is reflected in the incidence of clinical and subclinical mastitis within the US: approximately 20–25 cases per 100 cows per year. Clinical mastitis occurs in all dairy herds, even those that are well-managed [2]. Therefore, there is a tremendous need for safe and effective antimicrobials for treating this disease.

The ability to predict drug solubility in milk is important both for human and veterinary medicine. In terms of its veterinary application, a number of compounds are approved for the treatment of bovine mastitis via intramammary infusion (IMM). This method of drug administration typically involves the insertion of a syringe tip into the teat canal whereupon the contents of the syringe are expelled into the infected quarter. Subsequently, the quarter is gently massaged to facilitate distribution of the medication (e.g., see FOI for NADA #108-114; cephapirin benzathine gel). When administered into the bovine mammary gland, the drug acts locally within the udder with minimal systemic absorption.

Given its unique characteristics in terms of proteins, fats, and ionic composition, it is not appropriate to evaluate the solubility of these products using typical aqueous buffer systems. Milk contains more than 20 proteins, along with a variety of fats and a wide range of ions [3]. Moreover, there is the potential for preferential binding to casein milk proteins, whey, or fat. In addition, we need to consider the structure of the fat globules as well as the structure and stability of casein micelles (which can be impacted by the mineral content of the milk). Therefore, we considered the possibility of developing a standardized milk matrix to support these assessments. However, given the problems we encountered during efforts to produce a synthetic milk medium (both in terms of the lack of availability of some of the necessary components and difficulty in replicating the physicochemical quality of milk from available components of the milk matrix), the possibility of developing some standardizable approach was not considered feasible. It was also unclear as to whether some of the differences that might be encountered across herds could alter conclusions pertaining to drug solubility [4,5]. Therefore, we explore the possibility of variations in milk composition by comparing readily available milk matrices (i.e., off-the-shelf milk products such as whole milk, skim milk, or reconstituted whole milk powder) versus that obtained from freshly obtained raw (unprocessed) bovine milk. Logistic challenges associated with obtaining the raw milk samples necessitated that only a single local farm was evaluated.

In our previous study, the aqueous solubility estimates of six drug compounds (amitriptyline, acetaminophen, dexamethasone, nifedipine, piroxicam and prednisone.) were similar when tested in whole milk, reconstituted whole milk, or raw bovine milk [6]. However, in that paper, the data were not evaluated from the perspective of its relevance to what might occur when drugs are administered by IMM. Therefore, in this manuscript we re-examine those data from the perspective of IMM infusions and compare those results to the solubility assessment of two approved salt forms of the beta-lactam antimicrobial, cephapirin. [ToMORROW (NADA 109-114), which contains cephapirin benzathine and ToDAY (NADA 97-222), which contains cephapirin sodium].

Cephapirin sodium is very soluble in water; insoluble in most organic solvents. The pH of a solution in water containing the equivalent of cephapirin 1% is between 6.5 and 8.5. In contrast, cephapirin benzathine is practically insoluble in water, in ether, and in toluene; freely soluble in alcohol; soluble in 0.1 N HCl [7].

Information pertaining to the two API’s are provided in Table 1.

**Table 1.** Basic information of cephapirin and the salt forms [8].

| Form | Formula | MW | CAS # | % Active<br>(based upon<br>MW) |
| --- | --- | --- | --- | --- |
| Cephapirin<br>(free form) | C<sub>17</sub>H<sub>17</sub>N<sub>3</sub>O<sub>6</sub>S<sub>2</sub> | 423.5 | 21593-23-7 | 100% |
| Cephapirin sodium<br>(1:1 salt) | C<sub>17</sub>H<sub>16</sub>N<sub>3</sub>NaO<sub>6</sub>S<sub>2</sub> | 445.4 | 24356-60-3 | 95.1% |
| Cephapirin<br>benzathine (2:1 salt) | C<sub>50</sub>H<sub>54</sub>N<sub>8</sub>O<sub>12</sub>S<sub>4</sub> | 1087.3 | 97468-37-6 | 77.9% |

## Materials and Methods

The materials and methodology used for the solubility assessment for the six API’s other than cephapirin sodium and cephapirin benzathine are provided in the manuscript by Li et al., 2022 [6]. Source information for the cephapirin sodium and benzathine is provided in Table 2.

**Table 2.**
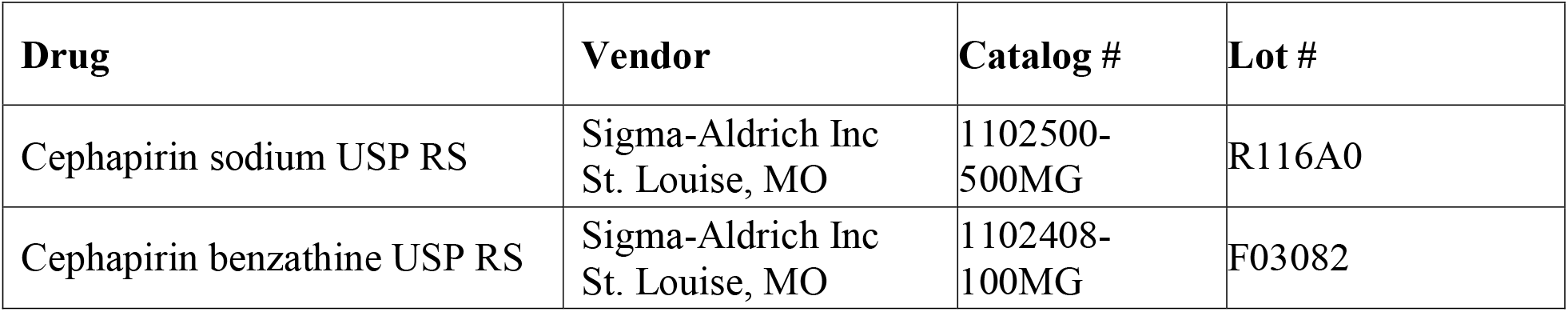
Cephapirin source information.

| Drug | Vendor | Catalog # | Lot # |
| --- | --- | --- | --- |
| Cephapirin sodium USP RS | Sigma-Aldrich Inc<br>St. Louise, MO | 1102500-500MG | R116A0 |
| Cephapirin benzathine USP RS | Sigma-Aldrich Inc<br>St. Louise, MO | 1102408-100MG | F03082 |

The source information for the vehicles used for the solubility testing are provided in Table 3. The milk powder was reconstituted with warm water (30–40°C) at 13% w/v. All vehicles were stored under refrigeration (2–8°C) until use. Raw milk and reconstituted whole milk were used within 24 hours. Skim and whole milk were used within expiration dates.

**Table 3.** Vehicle information.

| Vehicle | Vendor/Manufacturer |
| --- | --- |
| Skim Milk | Wegmans Food Markets, Rochester, NY |
| Whole Milk | Wegmans Food Markets, Rochester, NY |
| Whole Milk Powder (Nestle<br>Nido® brand) | Walmart, Rochester, NY |
| NIST Whole Milk Powder<br>(SRM 1549a) | National institute of standards and technology, Gaithersburg, MD |
| Raw Milk | Craft Dairy Farm, Williamson, NY |

Two types of syringe filters were purchased from Phenomenex (Torrance, CA) and used for solubility sample treatment prior to HPLC analysis. These included Phenex-GF 1.2 μm filters (Cat# AF0-8515-12) and Phenex-RC 0.45 μm filters (Cat# AF0-3103-52).

A Milli-Q Direct 8 system from Millipore Sigma (Burlington, MA) was used to produce purified water. Acetonitrile and trifluoroacetic acid (TFA), both HPLC grades, were purchased from Thermo Fisher Scientific (Waltham, MA).

### Physicochemical characterization of milk vehicles

The milk vehicles were characterized by pH, osmolality, viscosity, and globule size distribution. A 10 mM sodium phosphate buffer pH 6.8 was also included as a control.

#### pH

A Mettler-Toledo (Columbus, OH) Seven Easy model pH meter was used with a gel-filled pencil-thin pH electrode from Thermo Fisher Scientific (Waltham, MA). All vehicle samples were equilibrated to room temperature prior to the pH measurement.

#### Osmolality

A μOsmette Model 5004 osmometer from Precision Systems (Natick, MA) was used for osmolality measurement, which operates based on the freezing point depression method. For each measurement, 50 μL of the vehicle was placed in the sample tube and lowered into the freezing chamber. After the solenoid-induced pulse freezing, the liberated heat of fusion was related by the microprocessor to the freezing point of the sample, and the osmolality was automatically calculated and displayed.

#### Viscosity

A DHR-2 rheometer with a Peltier temperature-controlled cup and a double-gap concentric cylinder measurement system from TA Instrument (New Castle, DE) was used for viscosity measurements at 25°C. For each measurement, 12 g of the vehicle was placed in the cup, and the shear rate was scanned from 0.001 to 1 sec^−1^ on a log-scale with 5 points/decade. The method was set to determine the steady-state viscosity result with a maximum equilibration time of 480 sec at each point.

#### Globule size distribution

A LA-960 laser scattering particle size analyzer from Horiba (Piscataway, NJ) was used to analyze the globule size distribution of the milk related vehicles at room temperature. A refractive index of 1.45 was used for the oil phase and 1.333 for the aqueous phase. The milk was added dropwise to deaerated water to achieve 90 – 95% transmission for the red laser. A circulation speed of 2 was used with no sonication, and a data acquisition of 10,000 per read.

### Solubility assessment and sample treatment for HPLC analysis

Solubility assessment was performed in all milk vehicles listed above. A 10 mM sodium phosphate buffer pH 6.8 was also included as a control. For each solubility sample, an excess amount of drug powder was weighed in a 20-mL glass vial. A suitable volume of the vehicle, pre-equilibrated to the desired temperature, was added to the drug powder. The samples were stirred for 60 minutes at the same desired temperature. Upon completion, approximately 2 mL of the solubility sample was passed through the Phenex-GF 1.2 μm filter to remove the excess drug solid. As noted in manuscript by Li et al., 2022 [6], that pore size does not filter out the milk proteins present as whey or casein micelles, or the drug entrapped within fat globules. An aliquot of 0.5 mL filtrate was accurately transferred to a microcentrifuge tube followed by the addition of 1 mL extraction solvent. The extraction solvent was 0.1% TFA in acetonitrile (mobile phase channel B). The mixture was then centrifuged at 10,000 rpm, and about 1 mL of the supernatant was passed through a Phenex-RC 0.45 μm filter. The filtrate was collected in an HPLC autosampler vial and analyzed by the HPLC method described below.

Solubility testing was conducted at room temperature (RT) = 20-25°C, or at a temperature comparable to the rectal temperature the healthy adult cow, 38°C [9].

### HPLC analysis

A gradient HPLC method, described in Table 4, was used to analyze the concentration of all drug solubility samples after treatment.

**Table 4.** HPLC Method Parameters.

|  |  |
| --- | --- |
| HPLC System | Shimadzu Model# LC-2010AHT |
| Column | Phenomenex Kinetex C18, 150×4.6 mm, 5 µ, 100 Å (Phenomenex, Cat# 00F-4601-E0) |
| Column Temperature | 40 °C |
| Mobile Phase | Channel A: Water with 0.1% v/v TFA<br>Channel B: Acetonitrile with 0.1% v/v TFA<br>Linear gradient program: 5% to 60% Channel B in 10 minutes with 5 minutes of re-equilibration time.<br>Flow rate: 1.0 mL/min |
| UV Detection | 260 nm |
| Inject Volume | 10 µL |

## Results

When the highly soluble cephapirin sodium was added to the various milk samples, the color of the sample changes from white to deep yellow, even when tested at RT (Figure 1). However, increasing the temperature enhanced solubilization. This is illustrated by the increase in sample clarity in raw and reconstituted whole milk when the temperature is elevated from RT to 38°C (Figure 2).

**Figure 1:**
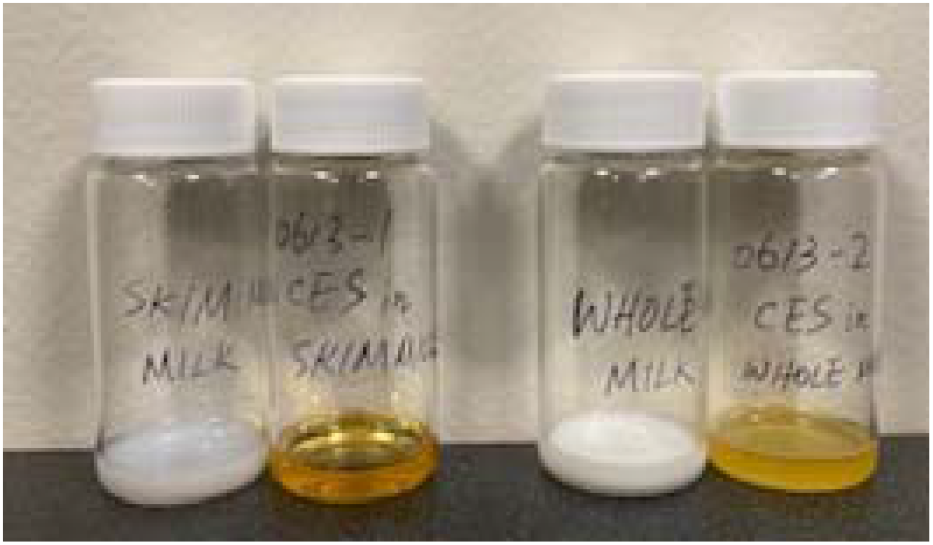
Photos of cephapirin sodium (CES) samples in various milk media at RT. Left to right: skim milk control, CES in skim milk, whole milk control, CES in whole milk.

**Figure 2:**
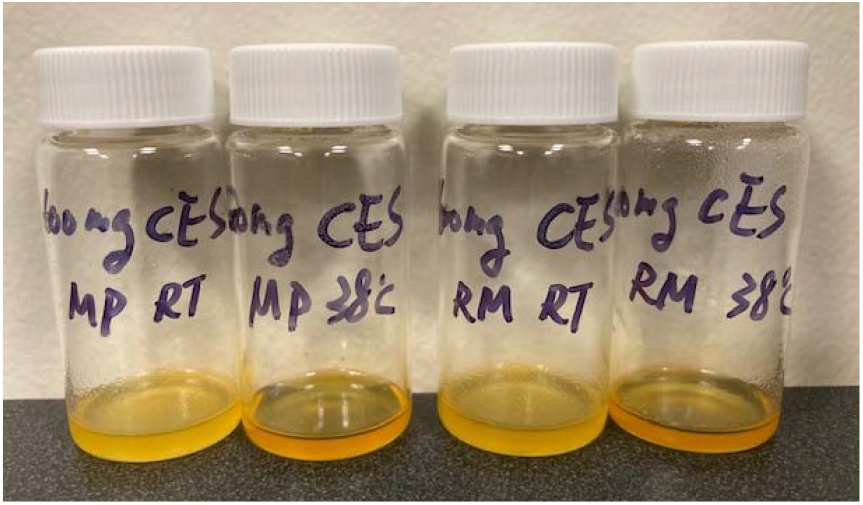
Photos of cephapirin sodium (CES) samples in various milk media at RT or 38°C. Left to right: CES in recon milk powder RT, recon milk powder 38°C, raw milk RT, and raw milk.

### *Physicochemical properties of milk media* (Table 5)

The pH values of all milk vehicles ranged from 6.6 to 6.8 (Table 5). With the except of the reconstituted whole milk powder, all other milk forms exhibited similar osmolality. The reason for the lower osmolality of the reconstituted Nestle and NIST whole milk powder is speculated to be attributable to an incomplete dissociation of the ingredients in water. All products except the reconstituted whole milk powder from NIST exhibited a viscosity of <5 mPas. This disparity is consistent with the larger globule diameter of the NIST product, being about 10-fold greater than that of raw milk. The larger globule size of the raw milk as compared to whole milk is not surprising given that the whole milk was homogenized. Similarly, the skim milk is effectively devoid of milk fat, rending the globule sizes about 5.3-fold less than that of the whole milk. While most of the data in this table was presented in the prior publication, it is listed again here for comparison with the new data generated with the NIST powder.

**Table 5.** Properties of Milk Related Vehicles.

| Vehicle | pH | Osmolality<br>(mOsm/kg) | Viscosity at 1 sec<sup>-1</sup><br>(mPas) | Globule size D<sub>50</sub><br>(µm) |
| --- | --- | --- | --- | --- |
| Skim milk | 6.65 ± 0.05 | 282 ± 1 | < 5 | 0.136 |
| Whole milk | 6.68 ± 0.04 | 277 ± 1 | < 5 | 0.726 |
| Raw milk | 6.70 ± 0.01 | 284 ± 2 | < 5 | 3.61 |
| Reconstituted whole<br>milk powder (Nestle) | 6.72 ± 0.03 | 234 ± 2 | < 5 | 0.512 |
| Reconstituted whole<br>milk powder (NIST) | 6.61±0.01 | 242±1 | 35 | 33.7 |

The NIST powder was included in this study by virtue of its standard chemical composition. Given the large globule size and high viscosity, it did not provide the physical properties of raw milk upon reconstitution and was therefore excluded from any further analysis. A potential reason for these differences may be the amount and composition of fats in the NIST product (as described in SRM 1549a) versus what has been reported in whole milk (e.g., that reported by the USDA [3]).

### Solubility

With the exception of the pH 6.8 aqueous buffer, the solubility of cephapirin sodium exceeded 350 mg/mL in the tested media. This is consistent with its classification of freely soluble. In contrast, markedly lower solubility was observed for the benzathine salt. Interestingly, the solubility in raw milk was somewhat lower than that observed in whole milk or whole milk powder but rather was more closely aligned with that of the skim milk. Nevertheless, across all media, the USP solubility classification of very slightly soluble remained the same (Table 6).

**Table 6.**
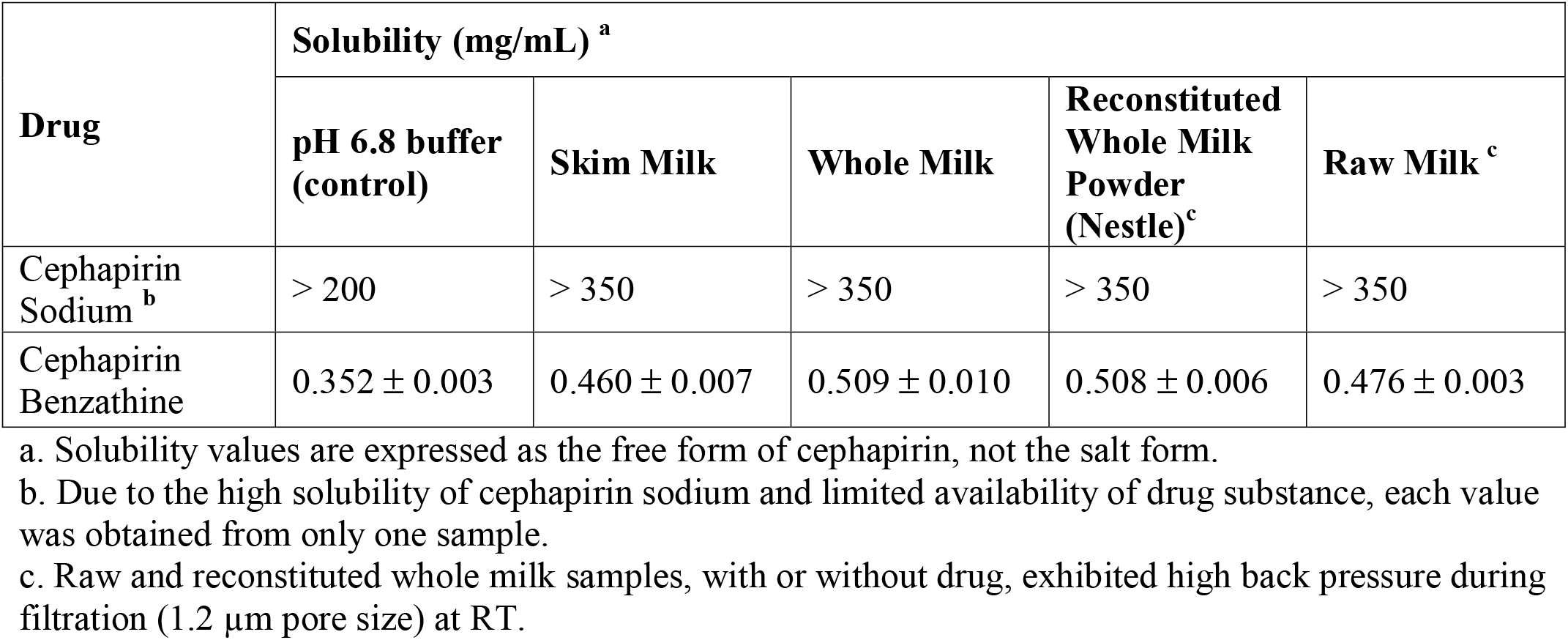
Solubility of cephapirin in selected milk vehicles at room temperature (n=3)

Since this study was directed specifically for drugs used for IMM infusion, we repeated the assessment of solubility using raw or reconstituted whole milk maintained at 38°C. Due to limited quantities of the API source, only two vehicles were tested at 38°C. This temperature is within the range of normal body temperatures for an adult cow (37.8 – 39.2°C) [9]. We also used two lots of milk powder to determine the potential for interlot variation. The results of these assessments are shown in Table 7.

**Table 7.** Solubility of cephapirin in selected milk vehicles at 38°C and nifedipine at RT or 38°C (n=3)

| Drug | Solubility (mg/mL) <sup>a</sup> |  |  |
| --- | --- | --- | --- |
|  | Reconstituted Whole Milk Powder (Nestle) |  | Raw Milk |
| Nifedipine RT | 0.0334 ± 0.0067 |  | 0.0120 ± 0.0014 |
| Nifedipine 38°C | 0.0947 ± 0.0143 |  | 0.0396 ± 0.0043 |
| Cephapirin Sodium <sup>b</sup> | > 500 |  | > 500 |
| Cephapirin Benzathine<sup>c</sup> | Lot 1 | Lot 2 | 0.706 ± 0.021 |
|  | 0.740 ± 0.005 | 0.762 ± 0.028 |  |

a. Solubility values are expressed as the free form of cephapirin, not the salt form.
b. Due to the high solubility of cephapirin sodium and limited availability of drug substance, each value was obtained from only one sample.
c. Three runs per lot of reconstituted whole milk.

In general, although its USP solubility classification [10] was not influenced by temperature, the use of higher temperatures increased the mg/mL of cephapirin dissolved from both salt forms, with the ratio of solubility results at 38 °C/RT ranging between 1.42 – 1.50. Therefore, the temperature-associated increase in cephapirin solubility was comparable in either reconstituted whole milk or raw milk. Similar outcomes were observed with nifedipine where the solubilized ratios at 38 °C/RT were 2.8 and 3.3 in whole and raw milk, respectively [6].

In terms of the relationship to cephapirin solubility in raw milk, less than a 10% difference was seen across all milk media. Only the aqueous buffer failed to provide quantitatively similar results. The ratio of mean solubility values across all media tested RT or 38°C) is provided in Figure 3.

**Figure 3:**
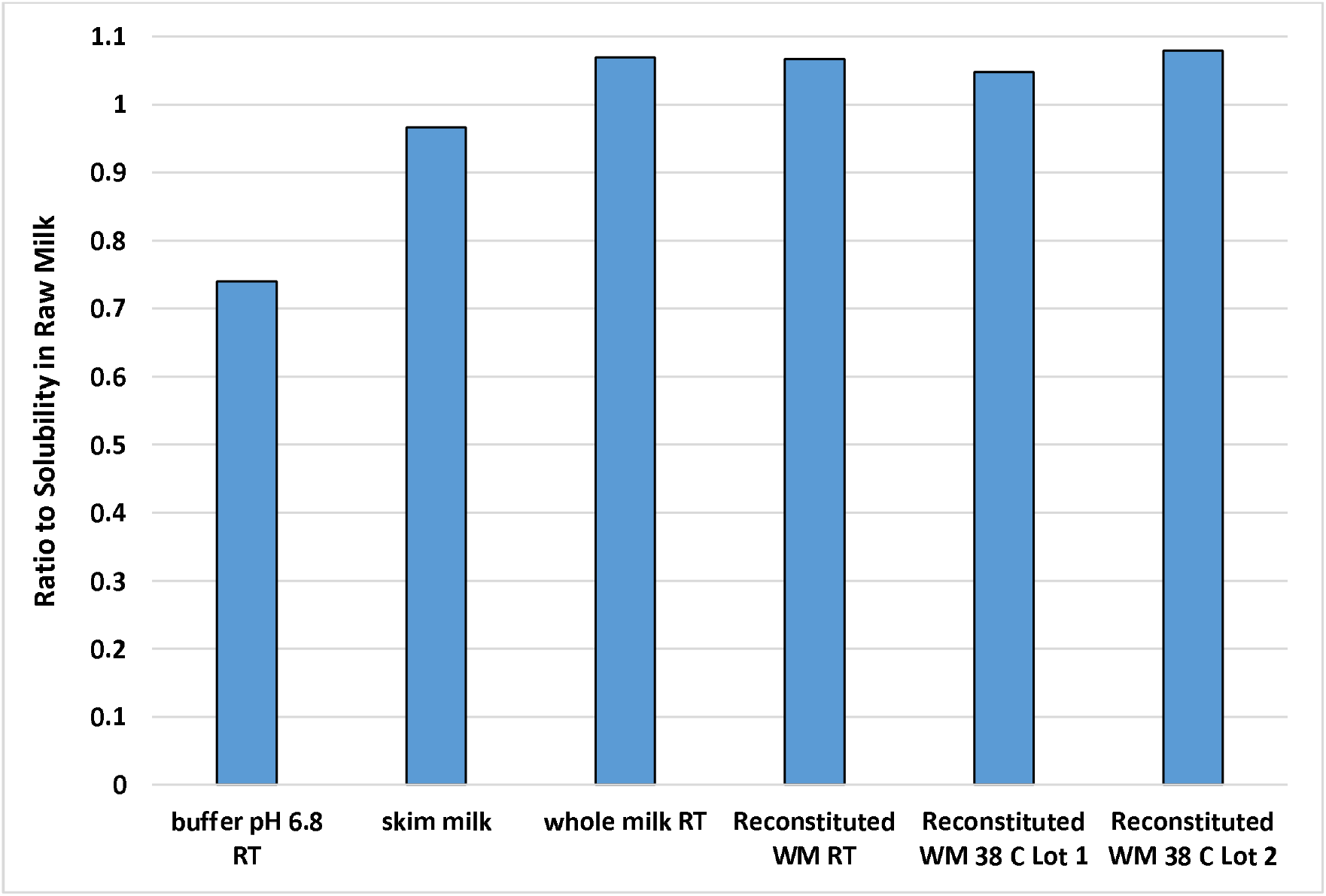
Ratio of cephapirin benzathine solubility across the three milk media and at RT versus 38°C.

One important difference between observations at 38°C versus RT was that the raw and reconstituted whole milk samples at RT, with or without drug, exhibited elevated back pressure during filtration (1.2 μm pore size). This problem was not encountered when the test was conducted at the bovine body temperature.

## Discussion

The goal of this study was to identify readily available media that could be used to predict the solubility of compounds when administered via infusion into the bovine mammary gland. To meet this goal, both previously examined model drugs and two products approved by the FDA for IMM infusion (cephapirin sodium and cephapirin benzathine) were evaluated in aqueous buffer, skim milk, whole milk, reconstituted whole milk, and raw bovine milk. Along with studying the effects of media, we also examined potential experimental challenges that could arise as a function of this assessment.

Filtration problems encountered with reconstituted whole milk and raw milk at room temperature were not seen when testing was conducted at 38°C. While this may have had some contribution to the temperature-associated differences in drug solubility in raw versus reconstituted whole milk, it is unlikely to have had a substantial impact since the solubilities in raw milk, reconstituted whole milk, and whole milk were similar under room temperature conditions (with whole milk not having the same filtration challenges).

Regarding the testing at RT versus 38°C, we find that the higher temperature is associated with greater solubility estimates. An even greater difference was seen for nifedipine in the previous study [6] where the solubility in raw milk and reconstituted whole milk increased approximately 3-fold. Since the increase in temperature is more physiologically relevant for the cow, the demonstrated increase in the solubility of active pharmaceutical ingredients across a range of physicochemical characteristics, along with the reduced problems associated with filtration pressure, leads us to conclude that when considering solubility testing of drugs intended for IMM infusion, testing should be conducted at 38°C.

Due to limitations with drug quantities, the day-to-day variability associated with the analytical method was not quantified. Therefore, it is unclear the degree to which this may have contributed to small differences seen between whole milk, raw milk, and lots of the reconstituted product. Nevertheless, given that the solubility values were typically similar across these three media, any bias in results due to inter-day variability should be minor.

With the exception of nifedipine, the solubility estimates of the model compounds provided by Li et al., 2022 [6] resulted in the same USP solubility classification across all media except that of the aqueous buffer. However, some differences were observed in the magnitude of drug solubility across the milk media used in this investigation. When considered relative to the solubility results obtained with raw bovine milk, that of whole milk was the one most consistently within ±10% that obtained in raw milk. All media failed to adequately reproduce the solubility of nifedipine, with whole milk and reconstituted whole milk having about 2–3-fold higher solubility than that in raw milk. In addition, skim milk failed to adequately reproduce the solubility piroxicam, a compound that for which similar solubility was observed in whole milk, reconstituted whole milk, and raw milk. The ratio of solubility estimates in the various media versus that in raw milk is provided in Table 8.

**Table 8:** Ratio of solubility estimate across compounds in the various media versus raw milk in RT (data from Li et al., 2022 [6]).

|  | pH 6.8<br>buffer | Skim<br>milk | Whole<br>milk | Reconstituted<br>whole milk |
| --- | --- | --- | --- | --- |
| acetaminophen | 1.05 | 1.08 | 1.05 | 1.01 |
| dexamethasone | **0.64** | 1.04 | 1.08 | 1.16 |
| nifedipine RT | **0.43** | **1.53** | **2.53** | **2.78** |
| piroxicam | **0.74** | **0.67** | 0.87 | 0.88 |
| prednisolone | **0.33** | 1.04 | 1.02 | 1.00 |

An additional uncertainty was whether drug solubility in homogenized whole milk, would be comparable to that in raw milk. In particular, the raw milk contains milk fat globules, where the lipids are contained within a membrane consisting of proteins, cholesterol, and phospholipids. In contrast, the homogenization process alters the contents and geometry of the milk fat globules, with the nature of the change varying as a function of method pressure and temperature [11]. Whether the availability of lipids within the natural membrane structure impacts the solubilization of lipophilic compounds has yet to be determined. However, from the data generated in this investigation and that of the model compounds [6], it can be concluded that any difference in the solubility of drugs in whole versus raw bovine milk will be relatively small.

The results obtained with nifedipine are particularly interesting since it is a weak acid (pKa 13) that is insoluble in aqueous (~10 μg/mL in water at 37°C), soluble in organic solvents, has a log K_OW_ of 2.2 and is known to present problems associated with in vitro dissolution [12,13,14]. Therefore, we expected that its solubility in whole milk would provide a far better reflection of raw milk as compared to that seen with skim milk. However, this was not the case, as shown in Table 8. In part this may be a function of the homogenization process allowing for greater dispersion of the nifedipine within the homogenized whole milk (i.e., enhancing its solubility) as compared to that associated with the raw milk where the fat is present as distinct globules or in skim milk. Nevertheless, this does not explain why the solubility in skim milk was still about 50% higher than that seen with raw milk. That this was not associated simply with the aqueous phase of the milk is supported by the lower solubility observed in buffer. Given the 92–98% plasma protein binding of nifedipine [15] the use of milk clearly provided an advantage over protein-free buffer systems in terms of solubilizing this API. Given that the protein contents of whole and skim milk are similar (15.4 g versus 16.5 g, respectively per 16 oz. glass) [16], there remains an uncertainty as to why it had a higher solubility as compared to that of raw milk. Clearly, further investigation is needed to understand these results more fully.

We know that the composition of raw and commercially available whole milk products can vary as a function of breed, diet, environment, and lactation stage [5]. Moreover, when administered into an inflamed and infected udder, further differences in milk composition are likely to exist [17]. Therefore, we cannot expect any singular medium to provide an absolute solubility value for an IMM dosage form. Correspondingly, the solubility assessment should strive to provide an assessment sufficient to determine a range of drug solubilities such that it can be characterized in accordance with the descriptive terms provided in USP 29 of USP characteristics [10].

## Conclusion

Since the model compounds, including the cephapirin benzathine, exhibited similar solubility in whole milk and in reconstituted whole milk, it appears that either store-bought whole milk or reconstituted whole milk could be used in lieu of raw milk to obtain an estimate of drug solubility within the bovine udder. Although we recognize that we did not evaluate the impact of factors that could influence the composition of bovine milk (as discussed in the introduction), the similarity in solubility estimates across these two sets of media suggest that normal variations that may occur in bovine milk under field conditions would not impact conclusions associated with the solubility of drugs intended for IMM. In that regard, because of its similarity to the solubility characteristics of raw milk, ready availability, and lesser challenges with respect to filtration, we conclude that testing solubility of API’s intended for IMM injection can be performed using shelf-ready whole milk. Lastly, given the impact of temperature on the solubility results, we recommend that all tests be conducted at 38°C.

## Ethics Statement

This work did not involve animal use.

## References

1. Ibrahim N. Review on mastitis and its economic effect. CJSR, 2017; 6(1): 13–22. https://doi:10.5829/idosi.cjsr.2017.13.22.

2. Cobirka M, Tancin V, and Slama P. Epidemiology and Classification of Mastitis. Animals (Basel). 2020;10(12):2212. https://doi:10.3390/ani10122212.

3. FoodData Central (usda.gov). https://fdc.nal.usda.gov/fdc-app.html#/food-details/746782/nutrients Accessed 09/21/2022.

4. Ogola H, Shitandi A, and Nanua J. Effect of mastitis on raw milk compositional quality. J Vet Sci. 2007;8(3):237–42. https://doi:10.4142/jvs.2007.8.3.237.

5. Linn JG. Factors affecting the composition of milk from dairy cows. National Research Council (US) Committee on Technological Options to Improve the Nutritional Attributes of Animal Products. Designing Foods: Animal Product Options in the Marketplace. Washington (DC): National Academies Press (US); 1988. Available from: https://www.ncbi.nlm.nih.gov/books/NBK218193/ Accessed 09/22/2022.

6. Li S, Gabriel JJ, Martinez MN, Longstaff DG, Coffey MJ, and Zhao, F. An exploratory study of a simple approach for evaluating drug solubility in milk related vehicles. BioRxiv, https://biorxiv.org/cgi/content/short/2022.10.27.514033v1

7. https://cdn.ymaws.com/www.aavpt.org/resource/resmgr/imported/cephapirin.pdf. Accessed 11/08/2022.

8. PubChem cephapirin. https://pubchem.ncbi.nlm.nih.gov/#query=cephapirin Accessed 09-28-2022.

9. Temperature reference ranges https://www.gla.ac.uk/t4/~vet/files/teaching/clinicalexam/examination/info/temperatures.html#:~:text=Normal%20Rectal%20Temperatures,101.5%2D103.5%C2%B0Fahrenheit%5D.

10. USP29: Description and Solubility. http://ftp.uspbpep.com/v29240/usp29nf24s0_desc-sol-2-5.html

11. Rui-Cano ME and Richter RL (1997). Effect of homogenization pressue on the milk fat globule membrane proteins. J Dairy Sci, 80:2731–2739. https://doi.org/10.3168/jds.S0022-0302(97)76235-0

12. Nifedipine | C17H18N2O6 - PubChem (nih.gov)

13. Friedrich H, Nada A, and Bodmeier R. Solid state and dissolution rate characterization of co-ground mixtures of nifedipine and hydrophilic carriers. Drug Dev Ind Pharm. 2005;31(8):719–28. https://doi:10.1080/03639040500216097.

14. Gajendran J, Krämer J, Shah VP, Langguth P, Polli J, Mehta M, Groot DW, Cristofoletti R, Abrahamsson B, and Dressman JB. Biowaiver Monographs for Immediate-Release Solid Oral Dosage Forms: Nifedipine. J Pharm Sci. 2015; 104(10):3289–98. https://doi:10.1002/jps.24560‥

15. Hazardous Substances Data Bank (HSDB): 7775 - PubChem (nih.gov)

16. Nutrition Comparison of Whole Milk vs Low-Fat Milk 2% vs Skim Milk (myfooddata.com)

17. Malek dos Reis, C.B., Barreiro, J.R., Mestieri, L. et al. Effect of somatic cell count and mastitis pathogens on milk composition in Gyr cows. BMC Vet Res. 2013; 9: 67. https://doi.org/10.1186/1746-6148-9-67

